# A Comparison of Weighted Stochastic Simulation Methods

**DOI:** 10.1101/2021.09.21.460158

**Authors:** Payton J Thomas, Mohammad Ahmadi, Hao Zheng, Chris J. Myers

## Abstract

Rare events are of particular interest in biology because rare biochemical events may be catastrophic to a biological system. To estimate the probability of rare events, several weighted stochastic simulation methods have been developed. Unfortunately, the robustness of these methods is questionable. Here, an analysis of weighted stochastic simulation methods is presented. The methods considered here fail to accomplish the task of rare event simulation, in general, suggesting that new methods are necessary to adequately study rare biological events.

## 2 INTRODUCTION

Despite occurring with low frequency, rare events can have devastating effects on biological systems. For example, rare biochemical events have been demonstrated to contribute to cancerous phenotypes by inactivating tumor-suppressing genes [1]. It is therefore important that computational methods be developed to analyze the probability of rare events.

Exact trajectories of biochemical reaction networks may be determined with *molecular dynamics*, wherein, given the initial position and momentum of each atom in the system, the complete state of the system can be determined at any time [4]. Unfortunately, such methods are computationally intractable for most systems. Instead, *stochastic chemical kinetics* (SCK) may be used to generate many potential trajectories for a system and approximate the probability of some event occurring [8].

Rare events can be problematic for stochastic simulation because the number of trajectories that must be generated to approximate the probability of a rare event may be computationally prohibitive. To address this issue, a variety of stochastic simulation algorithms have been developed that utilize *importance sampling* (IS) techniques to better estimate the probability of rare events [3, 6, 7]. In this abstract, three such algorithms are examined to determine how well they address the problem of rare event simulation.

The first algorithm that will be examined is the *weighted stochastic simulation algorithm* (wSSA) [6], which first applied IS techniques to biochemical network simulation. The second algorithm that will be examined is the *state-dependent biasing method for importance sampling* (swSSA) [7]. The third algorithm that will be examined is the *guided weighted stochastic simulation algorithm* (guided wSSA) [3].

## 3 RESULTS

The efficacy of each stochastic simulation method was tested on a six-reaction model of a biochemical futile cycle. This network is given as follows:

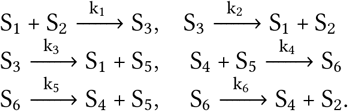

where

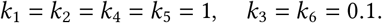

In the model, the initial state is

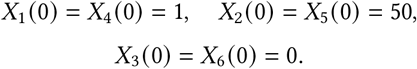

The rare event of interest is *X*_5_ → 40 within 100 time units, which is unlikely because the symmetry of the initial molecule counts and reaction rate constants will keep the system near the initial state with high probability. Futile cycles of this kind exist biologically in GTPase cycles, MAPK cascades, and glucose mobilization [2].

Rare events are difficult to simulate because the number of traditional SSA runs necessary to see a rare event of interest occur even once can be very high. Kuwahara and Mura solve this issue by increasing the likelihood of certain reactions occurring in simulation and decreasing the likelihood of others. Each run is then assigned a weight specific to the sequence of reactions that occurred such that the mean run weight is a sample estimator for the probability of the rare event of interest. This method requires that the user manually input the IS biasing parameters that are applied to each reaction.

In the small six-reaction example network, reaction three produces species five and reaction six consumes species five, so reaction three must be biased downward and reaction six must be biased upward. To this end, a single biasing parameter 0 < *δ* was introduced such that the rate of reaction three is multiplied by *δ* and the rate of reaction six is divided by *δ*. The performance of various magnitudes for *δ* is determined by comparing the true probability of a rare event *X*_5_ → 40 to the wSSA estimate after 10^2^ runs for 0 < *δ* ≤1.5 with increment 0.025 (Figure 1(a)).

**Figure 1:**
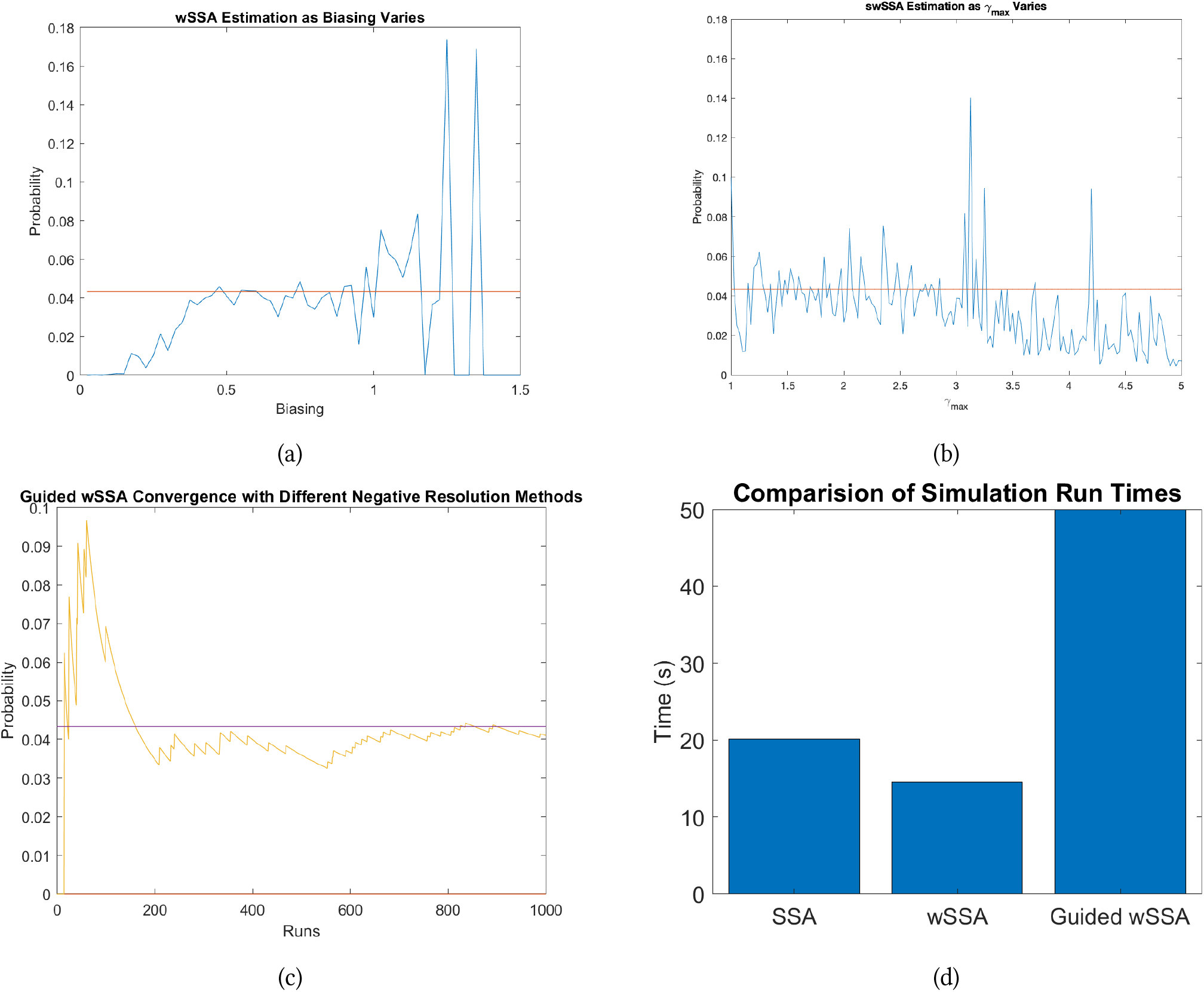
(a)True value of *P*_*t* ≤ 100_ (*X*_5_ → 40 |*x*_0_) (red) compared with the wSSA estimate at values of *δ* varying from 0.025 to 1.5 (blue). Note that *δ* = 1 corresponds to the traditional SSA, and *δ >* 1 corresponds to reciprocal weighting (decreases likelihood of reaching state of interest). (b)True value of *P*_*t* ≤ 100_ (*X*_5_ → 40 |*x*_0_) (red) compared with the swSSA estimate at values of *γ*_*max*_ varying from 1 to 5 (blue). (c) True value of *P*_*t* ≤100_ (*X*_5_ → 40 |*x*_0_) (purple) compared with the Guided wSSA estimate using each negative resolution method. Method A (yellow) and method C (blue) perform so similarly that method C is not visible. Method B fails to resolve negatives in general, and does not complete any runs. (d) The time to completion of 10^5^ runs of each algorithm is compared. The relative computational complexity of the Guided wSSA makes it much slower than other methods. The wSSA (with an ideal biasing parameter) performs faster than the SSA despite performing more calculations because a large proprotion of runs reach the state of interest before the total simulation time is reached.

As the species population changes throughout the course of simulation, relative propensity of each reaction changes too. The key insight presented in [7] is that forcing a fixed IS biasing factor in wSSA will result in a narrow range of values that factor can take to produce an accurate estimate. That is because the fixed biasing factor must adjust relative propensities appropriately for most of the possible values they take throughout the simulation. Also, a fixed biasing parameter will increase/decrease relative propensity of a reaction at a state where it already has a high/low probability of selection, resulting in lower accuracy. Therefore, [7] introduces a biasing factor which is a function of the relative propensity of the reaction it is adjusting at the current state. These functions are characterized by two sets of user inputs: (1) maximum amount of change allowed for each reaction, *γ*_*j*_ and (2) a threshold from which encouraging/discouraging reaction selection is stopped, *ρ*_0*j*_.

Figure 2 shows the results of estimating the probability of the rare event of interest on six reaction network. Again, *R*3 is set to be biased downward and *R*6 is set to be biased upward. Fixing *ρ*_0_ to be 0.6 for *R*6 and 0.2 for *R*3, and setting *γ*_3_ = *γ*_6_ = *γ*_*max*_, true probability of the rare event is compared to swSSA estimates with 1 ≤ *γ*_*max*_ ≤ 5 with increment 0.025 after 100 runs (Figure 1(b)).

To avoid reliance on user input and *a priori* knowledge of the system, Gillespie and Golightly calculate the conditioned expectation of reaction count over the remainder of the simulation for each reaction given that the rare state of interest is attained at the end of the simulation by assuming a constant reaction hazard and use that expectation to estimate an ideal amount of IS biasing [3, 5]. Unfortunately, the Guided wSSA may calculate a negative ideal biasing, and the resultant negative reaction rates cause errors in simulation. Inspection of the R code for the three example cases in Gillespie and Golightly reveals that a different method of dealing with these negatives is used in each case.

The Guided wSSA was ran using each of the three negative resolution methods with 10^3^ runs to compare the performance of each method (Figure 1(c)).

## 4 DISCUSSION

The wSSA requires the user to select which reactions should be encouraged and which reactions should be discouraged. Although such a task might seem trivial for very simple models, deep insight into underlying dynamics of the network is necessary for more complex models. Also, the biasing factor for those selected reactions must be specified prior to simulation. This is a tricky task, since these parameters can arbitrarily take any value greater than zero and it is by no means obvious what values will result in accurate estimates just by considering the model. Moreover, the accuracy of the estimate is highly sensitive to these values. Selecting non-optimal biasing factors can result in an estimate even less accurate than one produced by running the original SSA for the same number of simulations.

The swSSA suffers from the same issues. Reactions which are to be encouraged/discouraged should be specified by the user. Furthermore, for each of those reactions, the maximum amount of change allowed as well as a threshold from which encouragement/discouragement should be applied must be set prior to simulation, resulting in twice as many parameters as the wSSA. Like with the wSSA, the accuracy of this method is sensitive to these parameters, although the swSSA generally produces more accurate estimates and shows more robustness against a wider range of these parameters.

The Guided wSSA eliminates the need for specifying a set of reactions to bias and parameter(s) associated with each of those reactions as in the wSSA and swSSA. Since, in the Guided wSSA, matrices are inverted to automatically recognize a suitable biasing factor, this method is inherently slower then the SSA, wSSA, and swSSA in simulating trajectories. This additional computational effort may be justified if the run weights have a variance which is considerably smaller than those produced by other methods (as is the case with experiments discussed in [3]). Estimation of the probability of the rare event discussed in Section 3 on the six reaction network model using guided wSSA produces a far less accurate estimate than estimating that with wSSA while setting *δ* = 0.6. Running 200 simulations, it took guided wSSA 1.6 seconds to produce an estimate with the variance of 0.05 where it took wSSA 0.3 seconds to produce an estimate with the variance of 0.0015. The issue of complexity is demonstrated when the total runtimes of 10^5^ runs of each algorithm are compared (Figure 1(d)).

In summary, the original wSSA may achieve rapid convergence and lower variance than competing methods, but only with a narrow set of biasing parameters that cannot be reliably determined for an arbitrary system. The swSSA demonstrates broader robustness to biasing variation, but estimates with a high proportional error with few runs, lessening its advantage over the SSA. The guided wSSA solves the issue of biasing parameter determination, but has poor run-time performance and converges slower than the wSSA with optimal biasing.

## Acknowledgements

The authors of this work are supported by National Science Foundation Grant Nos. 1856740 and 1900542. Any opinions, findings, and conclusions or recommendations expressed in this material are those of the author(s) and do not necessarily reflect the views of the funding agencies.

## Notes

https://github.com/fluentverification/guided_proposals

### Competing Interest Statement

The authors have declared no competing interest.

## REFERENCES

[1] Esteller, M. Epigenetics in cancer. New England Journal of Medicine 358, 11 (2008), 1148–1159.

[2] Flomenbom, O., Velonia, K., Loos, D., Masuo, S., Cotlet, M., Engelborghs, Y., Hofkens, J., Rowan, A. E., Nolte, R. J., Van der Auweraer, M., et al. Stretched exponential decay and correlations in the catalytic activity of fluctuating single lipase molecules. Proceedings of the National Academy of Sciences 102, 7 (2005), 2368–2372.

[3] Gillespie, C. S., and Golightly, A. Guided proposals for efficient weighted stochastic simulation. The Journal of chemical physics 150, 22 (2019), 224103.

[4] Gillespie, D. Handbook of materials modeling, chapter 5.11, 2005.

[5] Golightly, A., and Wilkinson, D. J. Bayesian inference for markov jump processes with informative observations. Statistical applications in genetics and molecular biology 14, 2 (2015), 169–188.

[6] Kuwahara, H., and Mura, I. An efficient and exact stochastic simulation method to analyze rare events in biochemical systems. The Journal of chemical physics 129, 16 (2008), 10B619.

[7] Roh, M. K., Gillespie, D. T., and Petzold, L. R. State-dependent biasing method for importance sampling in the weighted stochastic simulation algorithm. The Journal of chemical physics 133, 17 (2010), 174106.

[8] Samoilov, M. S., and Arkin, A. P. Deviant effects in molecular reaction pathways. Nature biotechnology 24, 10 (2006), 1235–1240.

